# The archaeal division protein CdvB1 assembles into polymers that are depolymerized by CdvC

**DOI:** 10.1101/2021.10.07.463537

**Authors:** Alberto Blanch Jover, Nicola De Franceschi, Daphna Fenel, Winfried Weissenhorn, Cees Dekker

**Affiliations:** Department of Bionanoscience, Kavli Institute of Nanoscience Delft, Delft University of Technology, Delft, The Netherlands; Univ. Grenoble Alpes, CEA, CNRS, Institut de Biologie Structurale (IBS), 71, avenue des Martyrs, 38000 Grenoble, France

## Abstract

The Cdv proteins constitute the cell-division system of the Crenarchaea, in a protein machinery that is closely related to the ESCRT system of eukaryotes. The CdvB paralog CdvB1 is believed to play a major role in the constricting ring that is the central actor in cell division in the crenarchaea. Here, we present an *in vitro* study of purified CdvB1 from the crenarchaeon *M. sedula* with a combination of TEM imaging and biochemical assays. We show that CdvB1 self-assembles into filamentous polymers that are depolymerized by the action of the Vps4-homolog ATPase CdvC. Using liposome flotation assays, we show that CdvB1 binds to negatively charged lipid membranes and can be detached from the membrane by the action of CdvC. Interestingly, we find that the polymerization and the membrane binding are mutually exclusive properties of the protein. Our findings provide novel insight into one of the main components of the archaeal cell division machinery.

## Introduction

The Cdv system is the protein machinery responsible for cell division in the archaeal phylum of the Crenarchaeaota (*1*). Many components of this cell division machinery share a high degree of homology with the eukaryotic ESCRT machinery (*1, 2*) that is responsible for the cell division, vesicle budding, and multiple membrane-deforming processes in humans and yeast (*3*). This has led to the suggestion that the Cdv system is an evolutionary antique and simplified precursor of the eukaryotic ESCRT machinery (*4*), that may share the same mechanism at its core. While the eukaryotic protein complex is well studied, the complications of imaging live thermophilic cells such as the Crenarchaeaota, that live at ~85 °C, has long hindered a similarly fast growth in our understanding of the Cdv system.

Up until recently, most of our knowledge of the Cdv system was limited to CdvA, CdvB (an ESCRT-III homolog) and the AAA ATPase CdvC (Vps4 homolog), which are all found in the same operon (*1*). Basically, the CdvA protein was found to form a ring at the center of the cell together with CdvB (*1*). CdvA binds to the membrane in a spiral-like fashion (*5*) and acts as a membrane anchor for CdvB (*6*), as CdvB cannot bind to the membrane by itself (*7*). Subsequently, CdvC is recruited to this ring through the interaction of its MIT domain with the MIM2 domain of CdvB (*2*). In eukaryotes, several proteins of the ESCRT-III complex interact with the AAA ATPase Vps4 through this same MIM-MIT interaction (*8*), and the role of the ATPase is to detach the ESCRT-III proteins from the membrane and allow remodeling of the constricting filaments during the cytokinesis (*9*). Prompted by the homologies between the archaeal and eukaryotic proteins, and evidence for the presence of CdvC at the division ring during cytokinesis (*2*), it was thus believed that the role of CdvC in archaea is that of Vps4 in eukaryotes, namely removing CdvB from the membrane for its remodeling during the constriction of the membrane.

For a long time, the role of the CdvB paralogs (CdvB1, CdvB2 and CdvB3) was unclear. It was shown that *Sulfolobus* cells lacking CdvB were unable to grow, while cells lacking the CdvB paralogs were still viable, albeit with a lower growth rate or aberrant daughter cells (*10*), and therefore the CdvB1-3 paralogs were deemed non-essential for cell division. Recently however, developments in high-temperature microscopy techniques (*11, 12*) together with high-resolution imaging of fixed cells (*13*) shed valuable light onto how the CdvB paralogs participate in the cell division. It was shown that CdvB 1 and B2 are recruited to the CdvB ring right before the cytokinesis, whereupon CdvB disappears from the membrane, while CdvB1 and CdvB2 carry out the deformation of the membrane needed for the division of the cell (*11, 13*). Additionally, it was shown that *Sulfolobus* mutants without any CdvB2 undergo asymmetric cell divisions yielding differently sized daughter cells, while cells without CdvB 1 occasionally failed to divide, yielding multiploid cells with 2 genomes (*11*). All this changed the view of the basic mechanism of the Cdv system. The initial CdvA:CdvB ring at the center of the cell is now viewed as a non-contractile assembly ring that recruits CdvB1 and CdvB2 to the division site. Then, CdvC detaches CdvB from the ring, and CdvB1 and CdvB2 are left to deform the membrane and perform the division of the cell.

While this renewed model for the Cdv system arises, many questions remain. The constriction of the membrane requires a continuous and controlled disassembly of the contractile ring during the membrane deformation for successful scission and division of the cell (*13*). The division ring starts from a low-curvature conformation at the beginning of the division, and needs to proceed to an invagination of the membrane all the way down to the final step of scission. In this process, molecules that initially form the ring need to be removed to allow the final scission of the membrane to occur and avoid steric hindrance at the neck of the division site. It, however, remains unclear what drives these processes of depolymerization and constriction.

Since CdvB 1 also contains a MIM domain at the C-terminus, although slightly different from that of CdvB, it has been hypothesized that CdvC may also interact with CdvB1 through the MIT-MIM interaction, and CdvC may thus be responsible for the disassembly of the contractile ring (*13, 14*). In *S. islandicus,* yeast two-hybrid screenings showed interaction between CdvC and the CdvB paralogs (*15*), supporting the idea of its role in the disassembly of the contractile ring. However, there has so far been no experimental evidence that shows that this interaction leads to the depolymerization of structures formed by any of the CdvB paralogs. It is also unclear how the contractile ring stays bound to the cell membrane as the CdvB paralogs lack the wH domain that allows for the interaction with CdvA (*6*). Therefore, CdvB is presumed to act as a link between the membrane anchor CdvA, and CdvB1, which may start the recruitment of the contractile ring, but this raises the question of what links CdvB1 to the membrane after CdvB is gone. It has been proposed that the CdvA:CdvB ring does not fully disappear from the division site, but instead gets largely depolymerized, with a few proteins left behind which may be enough to hold the contractile ring in place (*14*). However, it has also been suggested that, in contrast to CdvB, the CdvB paralogs have a protein patch homologous to the membrane-binding domain of the human ESCRT-III CHMP3, with a certain degree of basicity to it, which could allow them to bind the membrane directly (*4*).

Here, we show how the CdvB paralog CdvB1 is able to polymerize on its own. We observe that heterologously expressed and purified CdvB1 proteins spontaneously self-assemble *in vitro* into filamentous structures. Furthermore, we show that CdvB1 filaments can get depolymerized by the action of CdvC, directly demonstrating both the presence of an interaction between these two proteins and the filament-remodeling activity of CdvC for the first time. Finally, we investigate the lipid-binding properties of CdvB1, and demonstrate that the ATPase activity of CdvC can detach CdvB1 from the lipid membrane.

## Results

### CdvB1 self assembles into polymeric filaments

CdvB1 and CdvC proteins from the archaeon *Metallosphaera sedula* (Figure 1A) were heterologously expressed in *E. coli* and purified (see Methods). To improve the handling and solubility of the protein, CdvB1 was fused to an MBP tag at the N-terminus of the protein, with an HRV 3C protease cleavage site in between the two. After the purification, the MBP was initially left on the protein, which largely impeded the polymerization of the protein, as can be seen by negative-staining TEM images (Figure 1B).

**Figure 1.**
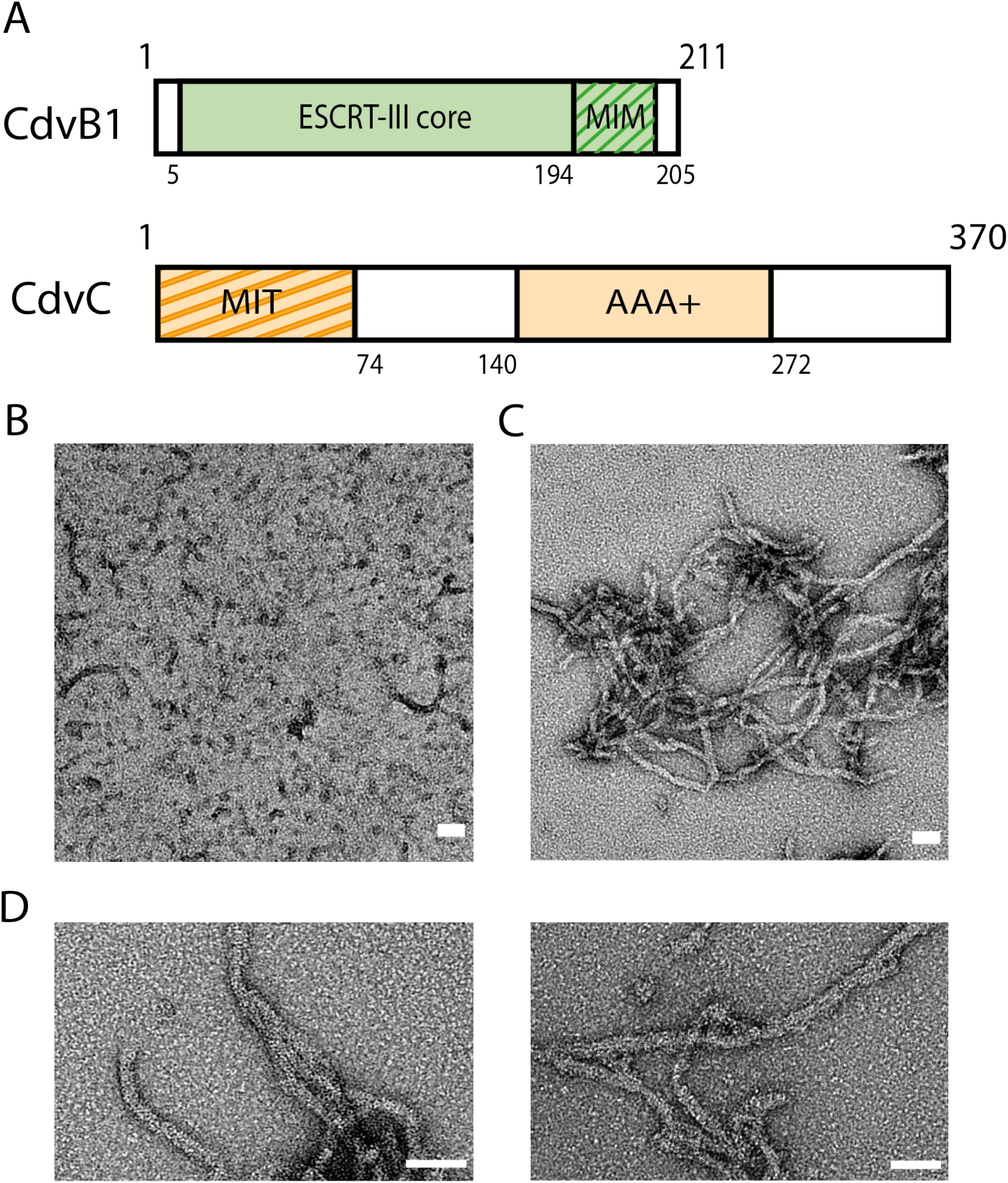
CdvB1 can polymerize into filamentous polymers. A. Schematic representation of CdvB1 and CdvC. B. Negative staining EM image of MBP-CdvB1 monomers and some short polymers. C. Negative staining EM image of the filamentous polymers formed by CdvB1 upon cleavage of the fused MBP. D. Closeup image of the CdvB1 polymers. All scale bars 50 nm.

It has been previously reported that CdvB, like many other ESCRT-III proteins, switches between an active and an inactive state when it comes to polymerization (*6, 16*). More specifically, CdvB contains a self-inhibiting domain that prevents it from polymerizing, while it spontaneously forms filaments when this domain is removed. We observed that this is not the case for CdvB1. Upon cleavage of the MBP by the 3C protease, CdvB1 spontaneously self-assembled into elongated filamentous polymers (Figure 1C), without the need of any other protein or the removal of any domain. The filaments have a defined width of about 14 ± 2 nm and a variety of different lengths, presenting an average length of 320 ±140 nm (Figure 1D). The filaments tended to stick to each other and thus form filamentous aggregates, making it difficult to properly measure the length when exceeding the 500 nm.

### CdvB1 polymers are disassembled by CdvC

Next, we studied whether these polymers of CdvB1 can get depolymerized by the action of CdvC. It has been hypothesized that CdvB1 can interact with CdvC, as some components of the ESCRT-III complex possess a MIM domain (MIT-interacting domain) at the C-terminus of their sequence, which interacts with the MIT domain of the CdvC/Vps4 ATPase. The sequences of these two interacting domains are highly conserved among species, and the CdvB 1 of various different crenarchaea exhibit, at the C-terminus of their sequence, a high degree of homology with the MIM2 domains of human’s CHMP6 or the yeast’s Snf7 (Figure 2A). Indeed, the MIM2 consensus sequence shows that prolines and hydrophobic amino acids are highly conserved at specific locations (*17*), and thus CdvB 1 has a high degree of sequence similarity to the human and yeast proteins.

**Figure 2.**
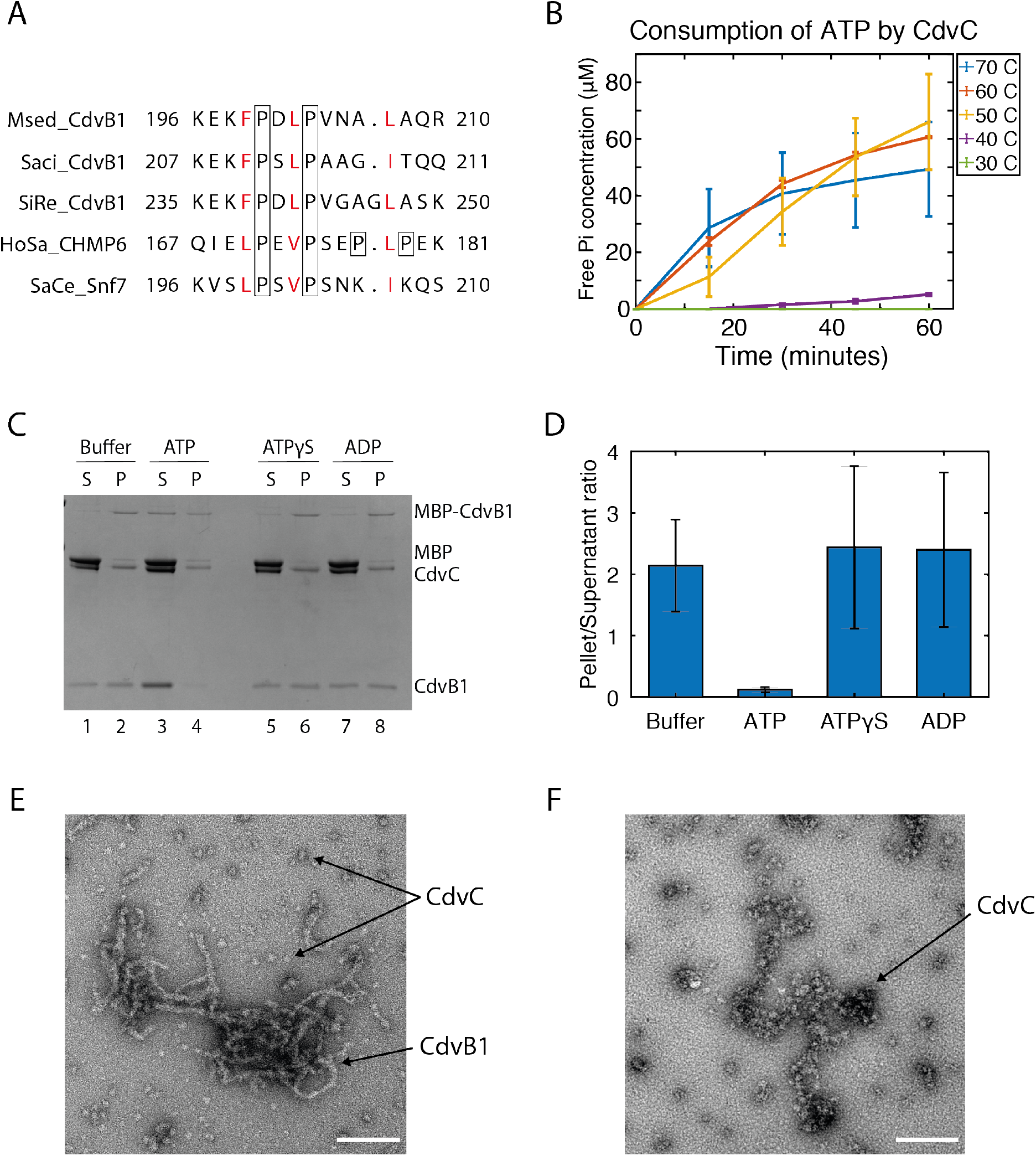
Filaments of CdvB1 are depolymerized by CdvB. A. Sequence alignment of regions of CdvB1 from *Metallosphaera sedula* (Msed2179), *Sulfolobus islandicus* (SiRe1200), and *Sulfolobus acidocaldarius* (Saci_0451), with the MIM2 domain of human ESCRT-III protein CHMP6 and yeast *Saccharomyces cerevisiae* Snf7. Conserved hydrophobic residues (red) and the prolines (boxed) are highlighted. B. ATP consumption by CdvC at different temperatures. C. SDS page analysis of a sedimentation assay where the depolymerization of CdvB1 filaments is assessed. D. Quantitative analysis of the Pellet-to-Supernatant ratios shown in the sedimentation assays (n=3). E. Filaments of CdvB1 with CdvC after incubation at 50°C but without ATP. F. EM image showing the disappearance of the CdvB1 filaments when mixed with CdvC, ATP and incubating at 50°C, leaving only aggregates of CdvC behind. All scale bars 50 nm.

The high temperature where the Crenarchaea live in their natural habitat posed an experimental challenge for testing the CdvB1-CdvC interaction. It had been previously reported that, *in vitro,* CdvC is enzymatically active at temperatures above 60°C (*6*). In our *in vitro* experiments, the CdvB1 polymers were broken down when incubated for prolonged times at temperatures above the 40°C, so we decided to verify whether a compromise between the ATPase activity of CdvC and the thermal stability of the CdvB1 polymers could be found. As shown in Figure 2B, CdvC did, as expected, not show any activity at temperatures up to 40°C. However, the protein did show a significant activity already at 50°C. At that temperature, together with the addition of Ficoll crowder, the CdvB1 polymers remained stable (Supplementary Figure 1), and hence, we chose this as our working temperature in the experiments.

To investigate if this interaction causes CdvC to disassemble CdvB 1 filaments, we first formed CdvB 1 polymers by cleaving off the MBP tag and allow them to polymerize. CdvC was then added to the samples, together with either ATP, ADP, non-hydrolysable ATP (ATPgS), or buffer without nucleotides (Buffer). These samples were incubated for 2 minutes at 50°C, to allow for the ATPase activity of CdvC, samples were centrifuged down at high speed, and pellet and supernatant were separately run on a gel. As can be seen from Figure 2C, polymerized CdvB 1 was forming pellets at the bottom during the centrifugation, while monomeric CdvB1 remained in the supernatant (Figure 2C, lines 1 and 2). However, when adding ATP and therefore allowing CdvC to act, the pelleted fraction virtually disappeared, and most of the protein was found to be in the monomeric state (Figure 2C, lines 3 and 4). The filament-to-monomer ratio of ~2 was the same for the samples where no hydrolysable nucleotides were added (ADP or ATPgS), whereas that ratio was drastically lowered to a value of 0.1 when CdvC could consume ATP (Figure 2D). This depolymerization did not occur when there was no CdvC present in the reaction (Supplementary Figure 1), indicating that the heating step did not disrupt the CdvB1 polymers, and it was indeed caused by CdvC.

We visualized the depolymerized filaments through negative staining EM. CdvB1 filaments were formed, CdvC was added and incubated at 50°C, as described above, and samples were imaged. When no hydrolysable nucleotide was added, CdvB1 filaments were observed in the sample, together with CdvC oligomers around them (Figure 2E). However, in samples where ATP had been added, the filaments of CdvB1 had vanished and only CdvC aggregates were observed (Figure 2F). Together with the evidences observed in the previous sedimentation assays, we thus conclude that the action of CdvC was responsible for the depolymerization of the CdvB1 filaments.

### CdvB1 binds negatively charged lipid membranes and can be detached by CdvC

Since the Cdv proteins are involved in remodeling membranes, it is of interest to study their membrane-binding properties. First, we studied if CdvB1 was able to directly bind lipid membranes. We used a liposome flotation assay, which, through a gradient of different concentrations of sucrose, allows distinguishing between the membrane-bound protein (that colocalizes with the liposomes), CdvB1 monomers (that stay in solution) and CdvB1 filaments (that precipitate to the bottom) (Figure 3A). We mixed MBP-CdvB1-Alexa488 with liposomes in a solution that contained the 3C protease. After incubation, we deposited the sample at the bottom of a 3-step sucrose gradient that we centrifuged at high speed. This resulted in 3 different fractions (Figure 3A): a top one where the liposomes were found (1), a middle one with monomeric protein (2) and a bottom one containing the filamentous CdvB1 (3). We tested liposomes made of DOPC + 0.1% Rhodamine-PE and a mixture of 70% DOPC + 30% DOPG + 0.1% Rhodamine-PE (percentages denote molar fractions) to examine the effect of the negative charges of the DOPG against the neutrality of DOPC. All the fractions of the gradient were analyzed by SDS PAGE where we imaged the fluorescence of both the lipids (red) and the CdvB1-Alexa488 (green). What we observed was that, as hypothesized, CdvB1 never bound to liposomes that were exclusively made of DOPC. However, when DOPG was present in the mixture, the CdvB1 protein showed clear binding to the liposomes (Figure 3B). This shows that it is not only CdvA that can bind lipid membranes, but that other components of the Cdv system can do so as well, similar to the way that different proteins of the ESCRT machinery in eukaryotes present different membrane-binding capabilities (*18*).

**Figure 3.**
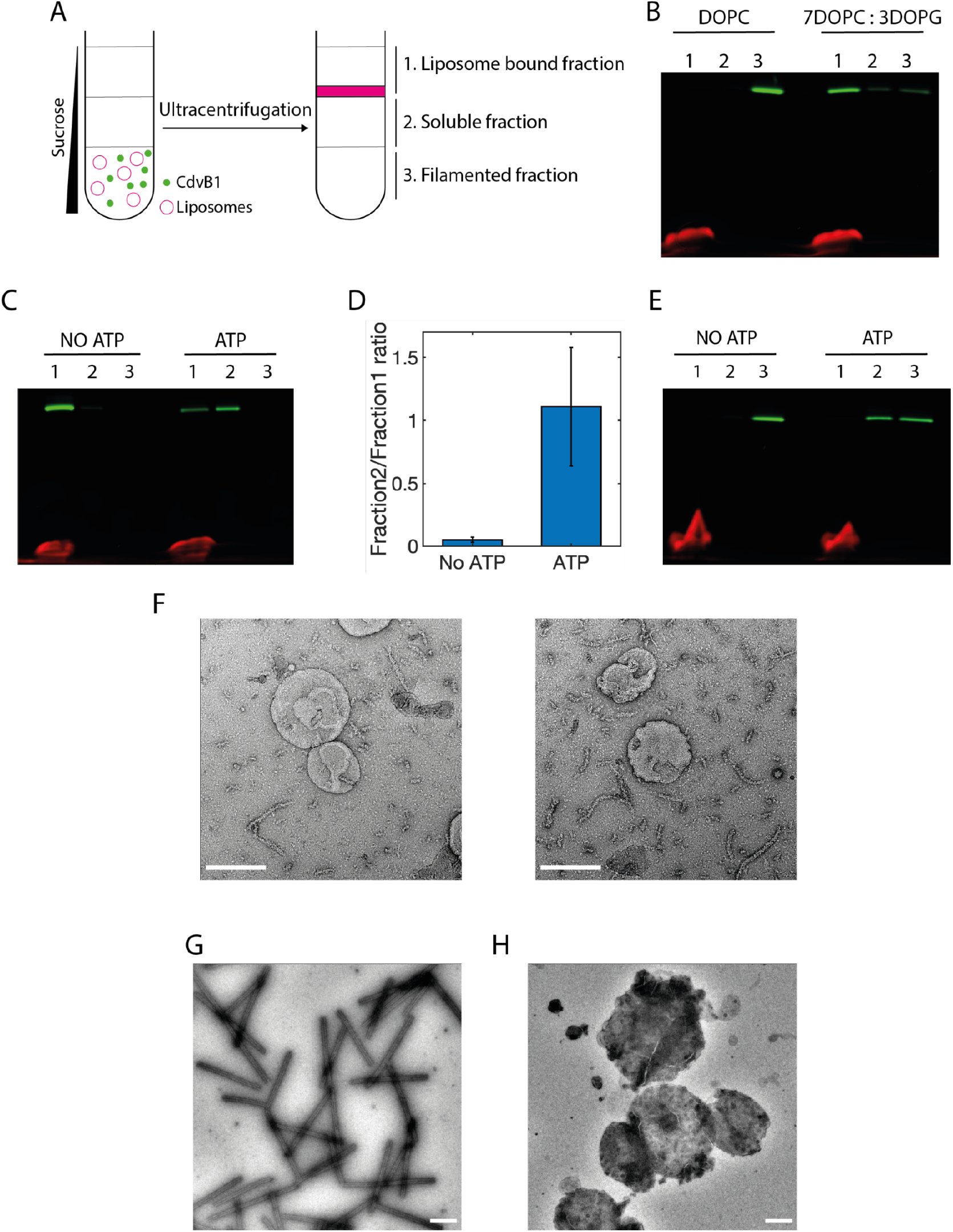
CdvB1 monomers bind to negatively charged lipid membranes but CdvB1 polymers do not. A. Schematic of the liposome flotation assay. B. SDS page gel of the liposome flotation assay showing CdvB1-Alexa488 in the liposome fraction only when the assay is performed in the presence of negatively charged lipids (30% DOPG). C. Liposome flotation showing that CdvB1 protein bound to liposomes is detached from the lipid membrane by CdvC. D. Quantitative analysis of the protein detachment from the membrane, showing the soluble-to-lipid-bound ratio (n=3). E. Liposome flotation assay showing that CdvB1 protein filaments do not bind to liposomes. The monomers resulting from their depolymerization do not bind either. F. Negative-staining EM image showing that the CdvB1 protein filaments do not interact with the membrane of the liposomes. Scale bars 200 nm. G. Negative-staining EM image of human ESCRT III proteins CHMP2A and CHMP3 co-polymerizing into tubelike structures. Scale bar 500 nm. H. Negative-staining EM image showing that CHMP3 does not form tube-like polymers when incubated with liposomes. Scale bar 500 nm.

In view of this, and the depolymerization of CdvB1 by CdvC that was described above, we tested whether CdvC was able to detach CdvB1 from the lipid membrane. For this we used the same liposome flotation assay where we bound the protein to negatively charged liposomes, subsequently added CdvC to the sample, incubated with ATP and Mg^2+^ for 10 minutes at 50°C, and then deposited the sample in the 3-step sucrose gradient. As can be seen in Figure 3C, we observed that the CdvB1 protein remained bound to the membrane in samples with no ATP. By contrast, samples containing ATP showed a big portion of the protein that disassembled from the liposomes to go into the soluble fraction (Figure 3C). On average, about half of the protein that was bound to the membrane depolymerized in solution in our experimental conditions (Figure 3D). When no CdvC was added to the reaction, this detachment was not observed (Supplementary Figure 2). These data show that, just like in eukaryotes, the same ATPase can interact with different proteins of the division system (in this case CdvB as well as CdvB1) and remove them from the lipid membrane (*19*).

This lipid-binding behavior was additionally tested for conditions where first CdvB1 filaments were allowed to form in the absence of liposomes, gently centrifuged to separate them from leftover monomers, then mixed with the liposomes, left to incubate for 1 hour at room temperature, and finally deposited and centrifuged in the 3-step sucrose gradient. Interestingly, we observed that, for these conditions, all of the CdvB1 was found in the bottom layer of the sucrose gradient, corresponding to the filamentous state, while no CdvB1 was found in the liposome fraction (Figure 3E). This shows that CdvB1 filaments, once formed, did not bind the membrane.

We then tried to see if depolymerization of the CdvB1 filaments with CdvC, would allow the newly solubilized CdvB1 to bind the lipids. For this, we formed CdvB1 filaments in the absence of lipids, mixed them with 7:3 DOPC:DOPG liposomes, CdvC, and ATP with Mg^2+^, and incubated for 10 minutes at 50°C. After the depolymerization reaction, we left the proteins with the liposomes rest at room temperature to interact for 1 hour, and then performed the 3-step sucrose gradient. We observed that neither the filaments nor the depolymerized CdvB1 showed binding to the liposomes (Figure 3E). This may suggest that CdvC is actually unfolding the monomers of CdvB1 in the process of depolymerizing the filaments, and that the resulting depolymerized proteins lack a functional folded structure. Negative-staining TEM images of MBP-CdvB1 incubated with vesicles that contained the 3C protease in solution showed filaments of CdvB1 that were lying next to the vesicles (Figure 3F). Hardly ever were these filaments found on top of the vesicles or attached to them, consistent with the results from the liposome flotation assay.

Given the similarities of the archaeal Cdv proteins and the eukaryotic ESCRT, we wanted to see if this duality between polymerization and membrane binding was present in both systems. Interestingly, a similar behavior was indeed observed for the CHMP2A and CHMP3, which are human homologs of CdvB1 (*4*). These proteins from the human ESCRT-III machinery are well known for their *in vitro* copolymerization into large helical structures that can be disassembled by the ATPase Vps4 (*20, 21*). These tube-like structures easily form when mixing MBP-CHMP2AΔC and CHMP3 at a molar ratio of 10:1 (Figure 3G). However, when trying to polymerize these tubes in the presence of liposomes (9:1 DOPC:PIP2), we found no tubular polymerization, as seen in Figure 3H. Instead, we observed that the protein remained bound to the surface of the liposomes, but it would never polymerize into helical tubes and bind to liposomes at the same time.

## Discussion

In this paper, we clarified a number of characteristics of the important but so far understudied Cdv protein CdvB1. We found that, *in vitro*, CdvB1 self assembles into filaments without the need of removing any inhibiting domain, like in many of its ESCRT-III homologs. Fusion to an MBP impeded activation and filament formation, which is a convenient way of controlling the polymerization when needed, facilitating its study in *in vitro* assays. We also showed how CdvB1 polymers are disassembled by CdvC, showing for the first time a direct proof of depolymerization of an archaeal ESCRT-III polymer by the action of the AAA ATPase CdvC. These findings strengthen the idea that the Cdv system can be considered as a relatively simpler version of the homologous eukaryotic ESCRT machinery, thus reinforcing evidence for a mechanistic common ground between the archaeal and eukaryotic cell division systems.

Our data show that CdvB1 has the ability to directly bind membranes with negatively charged lipids without the need of any anchoring proteins (such as CdvA), which contrasts previous findings for CdvB. Furthermore, CdvC was found to remove the protein from the membrane. This may explain how CdvB1 can stay attached to the membrane during the cell division after the removal of CdvB. This is consistent with the view that the initial CdvA:CdvB ring merely serves as a scaffold for the recruitment of CdvB 1 to the division site, whereupon the contractile ring can stay bound to the membrane by its own interaction with the lipids after the initial ring is removed. We know that the initial ring gets digested by the proteasome before the constriction of the cell (*13*), and it is likely that CdvC is the responsible for the removal of CdvB from the ring so it can be digested. Therefore, the same CdvC which first removes the initial scaffold ring, may in a later stage generate the depolymerization of the contractile ring that is needed for the division of the cell to occur (*13, 14*).

We also reported an interesting duality for CdvB1, as the polymerization of the protein and its membrane-binding capabilities were found to be mutually exclusive phenomena. After cleavage of the MBP tag, i.e., at the point where the protein is in its fully native conformation, one of two things can happen: either a CdvB1 protein finds the lipid membrane and binds to it, or it binds other CdvB1 proteins to polymerize into filaments that subsequently are not able to bind the lipid membrane. Interestingly, we showed a very similar behavior for the human ESCRT-III CHMP2A and CHMP3, that co-polymerize into large helical structures in the absence of lipids, but fail to do so when surrounded by liposomes. These data might be taken to suggest that upon polymerization, the membrane-binding patch of the proteins does not face the outside anymore, and thus loses its lipid-binding ability. However, this seems counter-intuitive regarding the function of the ESCRT-III proteins that form filamentous polymers that can be reshaped to deform the membrane. We suggest that instead, *in vivo*, CdvB1 is recruited in a monomeric state to the lipid membrane at the division site by CdvB, whereupon it may form filaments (see Figure 4). This would allow CdvB1 to bind to the membrane by its membrane-binding domain, where it may also recruit CdvB2 to the division site. There is a very high degree of conservation of positively charged amino acids between CdvB1 and CdvB2, which leads us to think that CdvB2 will likely bind to negatively charged lipid membranes as well. Therefore, we speculate that upon removal of CdvB from the membrane, with CdvB 1 and CdvB2 left to constrict the membrane, CdvB1 may polymerize into filaments that will be stabilized and kept bound to the membrane by CdvB2. The action of CdvC may then remodel the CdvB1 filaments whereupon the membrane shrinks, while CdvB2 keeps the ring in place. This idea is in line with previous findings (*11*) where mutants lacking CdvB 1 were less able to perform the fission of the membrane, indicating its role in membrane deformation, whereas mutants without CdvB2 would lead to asymmetric divisions, suggesting its guiding role.

**Figure 4.**
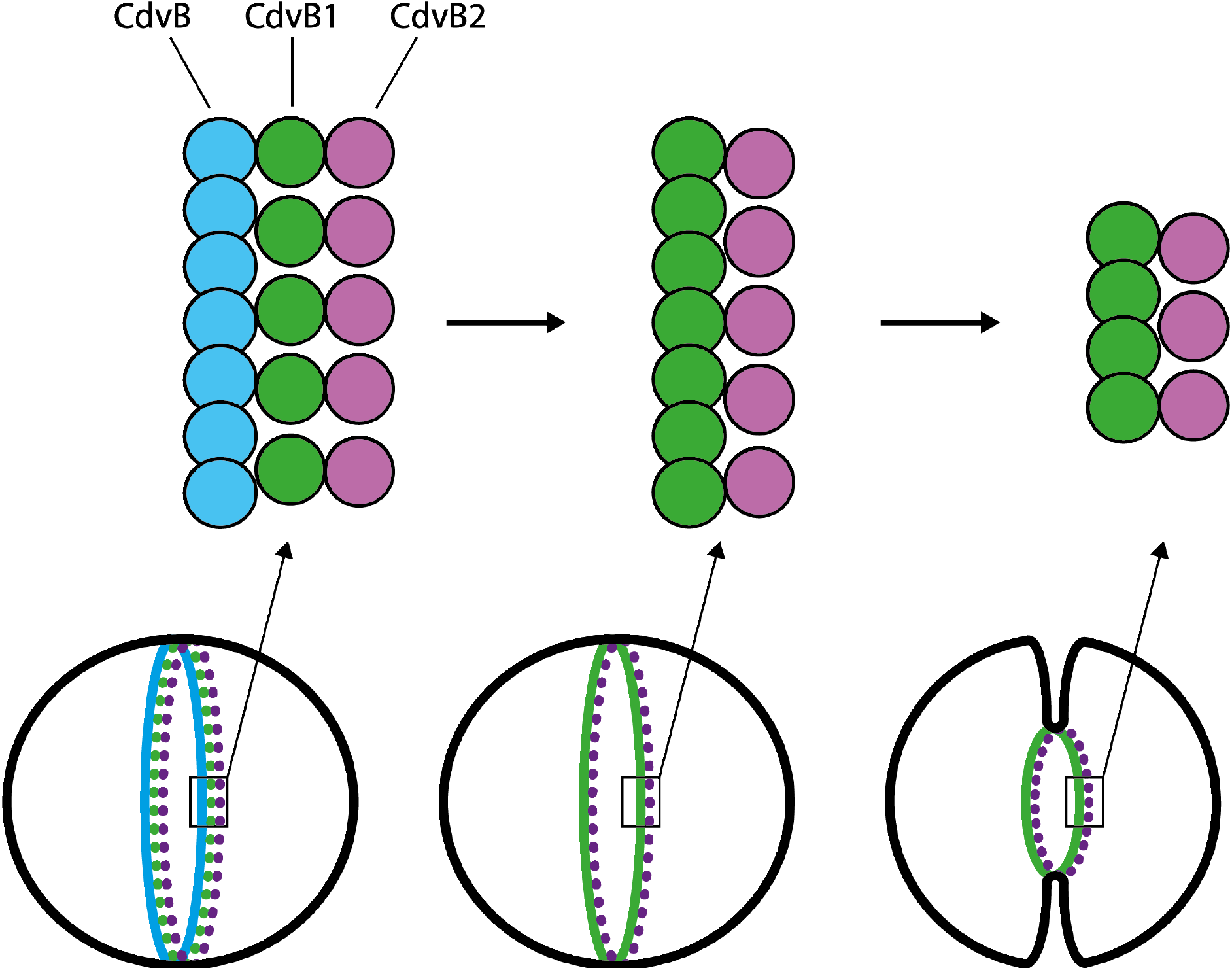
Proposed filament structure in crenarchaeal cell division. First CdvB forms a non-contractile ring, whereupon it recruits monomeric CdvB1 and CdvB2. Upon removal of CdvB, CdvB1 can polymerize while remaining in the right place thanks to the action of CdvB2, and jointly they constrict the membrane.

An evolutionary correlation of these phenomena seems to be implied in the clear *in vitro* polymerization versus membrane binding that we observed in both the human and archaeal ESCRT-III proteins. Previous studies showed the importance of conformational changes of ESCRT proteins bound to a lipid membrane, and how the different states of the protein, either bound or unbound, facilitated different interactions (*22*). A similar scenario might apply to the Cdv proteins where, depending on the interaction with the membrane, different polymerization states may occur. Future high-resolution studies of the proteins *in vivo* will help to better understand how the proteins arrange themselves at the division site to facilitate faithful cell division in the Crenarchaeaota.

## Acknowledgements

We thank Eli van der Sluis and Ashmiani van den Berg for assistance in protein purification and discussions; Patricia Renesto for the plasmid for CdvC; Yaron Caspi for discussions and early protein purification; and Jaco van der Torre for discussions and experimental assistance. We acknowledge funding support from the BaSyC program of NWO-OCW (024.003.019) and from the ERC Advanced Grant LoopingDNA (no. 883684). We acknowledge access to the platforms of the Grenoble Instruct-ERIC center (IBS and ISBG; UAR 3518 CNRS-CEA-UGA-EMBL) within the Grenoble Partnership for Structural Biology (PSB), with support from FRISBI (ANR-10-INBS-05-02) and GRAL, a project of the University Grenoble Alpes graduate school (Ecoles Universitaires de Recherche) CBH-EUR-GS (ANR-17-EURE-0003).

## Materials and Methods

### Plasmids

The gene for CdvB1 from *M. sedula* was obtained from the Gen Bank data base (Msed_2179), and was reverse translated using the EMBOSS Backtranseq tool, optimized for *E. coli* codon usage. To the resulting DNA sequence, a codon of a cysteine for fluorescent labelling was added at the N terminal of the protein, as well as Tobacco Etch Virus (TEV) and an HRV 3C proteases cutting sites. It was all ordered as a synthetic gene in a pMAL-c5x vector from Biomatik. The plasmid for CdvC was kindly provided by Patricia Renesto’s lab.

### Protein purification

The protein was expressed in a BL-21 *E. coli* strain. Cells were grown at 37°C in LB^amp^ medium to an OD of around 0.5. Expression was induced with a final concentration of 0.1 mM of IPTG for 4 hours. Cells were then harvested by centrifugation at 4500x g at 4°C for 12 minutes. The pellet was resuspended again in lysis buffer (50 mM Tris pH8.8, 50 mM NaCl, 50 μM TCEP, cOmplete™ Protease Inhibitor Cocktail (Roche)). Cells where lysed by French press, and the lysate was then centrifuged for 30 minutes at 45000 rpm in a Ti45 rotor (Beckman Coulter). Supernatant was then incubated rotating with 1ml of amylose resin (NEB) at 4°C for 2 hours. In a 4°C room, the lysate was then poured through a gravity chromatography column, then washed twice with 1 column volume of purification buffer (50mM Tris pH 8.8, 50mM NaCl, 50μM TCEP). The washed resin was incubated for 5 minutes with elution buffer (50mM Tris pH 8.8, 50mM NaCl, 50μM TCEP, 10mM maltose), and finally eluted the protein out using the same column. The protein was then concentrated down to a volume of 0.5 ml, and run through a Superdex™ 75 increase 10/300 GL size exclusion chromatography column mounted in an ÄKTA^TM^ Pure system. Sample ran with purification buffer, and purity of the eluted peaks was evaluated by a 12% SDS PAGE gel stained with Coomassie blue. After the resin column, a fraction of the protein was dialyzed into the same buffer composition but pH of 7.4, and labelled with Alexa488-maleimide. The rest of the purification from that point stayed the same. For CdvC, the purification was performed as indicated in (*6*).

### TEM imaging

MBP-CdvB1 at a final concentration of 1 μM was mixed with 0.1 μM of the 3C protease in buffer containing 50mM Tris pH 7.4, 50mM NaCl and, to allow filaments for form for at least 1 hour. The measurements of width and length of the filaments were extracted from 3 independent experiments. For samples with liposomes, lipids used were DOPC (1,2-dioleoyl-sn-glycero-3-phosphocholine), DOPG (1,2-dioleoyl-sn-glycero-3-phospho-(1’-rac-glycerol)), PIP2 (1,2-dioleoyl-sn-glycero-3-phospho-(1’-myo-inositol-4’,5’-bisphosphate)) and Rhodamine-PE (1,2-dioleoyl-sn-glycero-3-phosphoethanolamine-N-(lissamine rhodamine B sulfonyl)) and they were all purchased from Avanti Polar Lipids. LUVs of 400nm of 7:3 DOPC:DOPG were prepared by extruding the lipid mixture at a concentration of 5 mg/ml through a polycarbonate filter. LUVs were then diluted down to 0,5 mg/ml, mixed with 0.1 μM of 3C protease first, and then MBP-CdvB1 was added to a final concentration of 1 μM. It was all left to incubate at room temperature for at least 1 hour. The sample was then analysed by negative staining.

MBP-CHMP2AΔC and CHMP3 were purified as previously described (16). MBP-CHMP2AΔC at 10uM and CHMP3 at 1uM were mixed in 50mM Tris pH 7.4, 150mM NaCl, 1mM TCEP in the presence or absence of liposomes and incubated overnight. Liposomes were made with a mixture 9:1 of DOPC:PIP2. The sample was analysed by negative staining.

### ATPase activity assay

The ATPase activity assay was done using the Phosphate Assay Kit – PiColorLock™ from Abcam and performed according to the manufacturer’s guidelines. A final concentration of CdvC of 0.1 μM in buffer 50mM HEPES pH 7.5, 50mM NaCl and 5mM MgCl_2_ was tested with a concentration of 0.1 mM ATP at different temperatures (Room temperature, 30, 40, 50, 60, 70°C). The reaction was stopped at various time points (15, 30, 45 and 60 minutes) by submerging the samples in liquid nitrogen. Afterwards they were thawed, added the reagent of the kit and measured the absorbance at 630nm in a 96 well plate reader. A free phosphate standard curve was plotted to calculate the amount of phosphate released by the protease during the reaction. All experimental conditions were performed in triplicates and results were normalized to buffer with ATP under the same conditions without the ATPase.

### Sedimentation analysis of filament depolymerization

For the filament formation, 1μM of MBP-CdvB1 was mixed with 0.1 μM of 3C protease in buffer 50mM HEPES pH 7.4, 50mM NaCl, 5mM MgCl_2_, Ficoll 41.25 mg/ml. Incubated overnight at 4°C to guarantee full formation of the CdvB1 filaments. The next day, mix in 0.6 μM of CdvC and incubate for 30 minutes. For the depolymerization of the filaments, of the corresponding nucleotide was added to a final concentration of 1mM and then incubated at 50°C for 2 minutes in a thermocycler. After incubation, the sample was transferred to an ultracentrifuge tube and spun down in a Ti 42.2 rotor at 140,000 xg for 30 minutes at 4°C. After centrifugation, the supernatant was collected, the pellet was resuspended in the same volume, and they were analysed by SDS-PAGE stained with Coomassie.

### Liposome flotation assay for membrane binding

Protocol adapted from (*23*). Lipids were mixed to final ratios (mol:mol) of 99.9 DOPC: 0.1Rhodamine-PE or 69.9 DOPC: 30DOPC: 0.1Rhodamine-PE and evaporated in a glass vial to a final amount of 500 μg. They were later resuspended in 100 μl of buffer containing 50mM HEPES pH 7.5, 50 mM NaCl and 300 mM sucrose. The lipid film was hydrated for 1 hour and thoroughly vortexed to form small lipid vesicles. In an ultracentrifuge tube, 300 μg of the lipid vesicles were mixed with 3C protease and MBP-CdvB1-Alexa488, to a final concentration of 1,5 μM. Lipids and protein were left to incubate for 45 minutes, and then buffer with sucrose was mixed to obtain a bottom layer of 80 μl of 30% sucrose solution. Carefully, on top of it, a layer of the same volume of buffer with 25% sucrose was deposited, and another layer with 0% sucrose buffer on top of all. Then centrifuged at 200,000 xg at 21 °C for 30 minutes in a Ti 42.2 rotor. Then, all the different sucrose gradient layers were pipetted out and analysed by SDS-PAGE and imaged the fluorescence of protein and lipids with a GE Amersham™ Typhoon gel imager.

### Detachment of CdvB1 from the membrane by CdvC

Binding of CdvB 1 to the membrane was performed as described for the “Liposome Flotation Assay” but in a PCR tube instead of in the ultracentrifuge tube. CdvC was then added to a final concentration of 0.8 μM. Samples were the added 1:10 of either ATP stored at 10mM in buffer with 50mM HEPES pH 7.5, 50mM NaCl and 20mM MgCl_2_ or simply the ATP storage buffer. Samples were incubated for 10 minutes at 50°C in a thermocycler. After incubation, samples were moved to ultracentrifuge tubes and the sucrose gradient and subsequent centrifugation were performed as previously described.

### Assessment of the binding of pre-formed CdvB1 polymers

Polymers of CdvB1 were formed by cleavage of the MBP in absence of any lipids. After 1 hour, the filaments were spun down at 70,000xg for 20 minutes to separate the filaments from remaining monomers. Then, the protein was mixed with the same amount of liposomes and CdvC as previously described, and the monomer depolymerization reaction was carried out by adding ATP at 50°C for 10 minutes. After that, the sample was left at room temperature to allow any binding of the proteins for 45 minutes, and then the sucrose gradient and subsequent centrifugation were performed as previously described.

## Supplementary

**Supplementary Figure 1.**
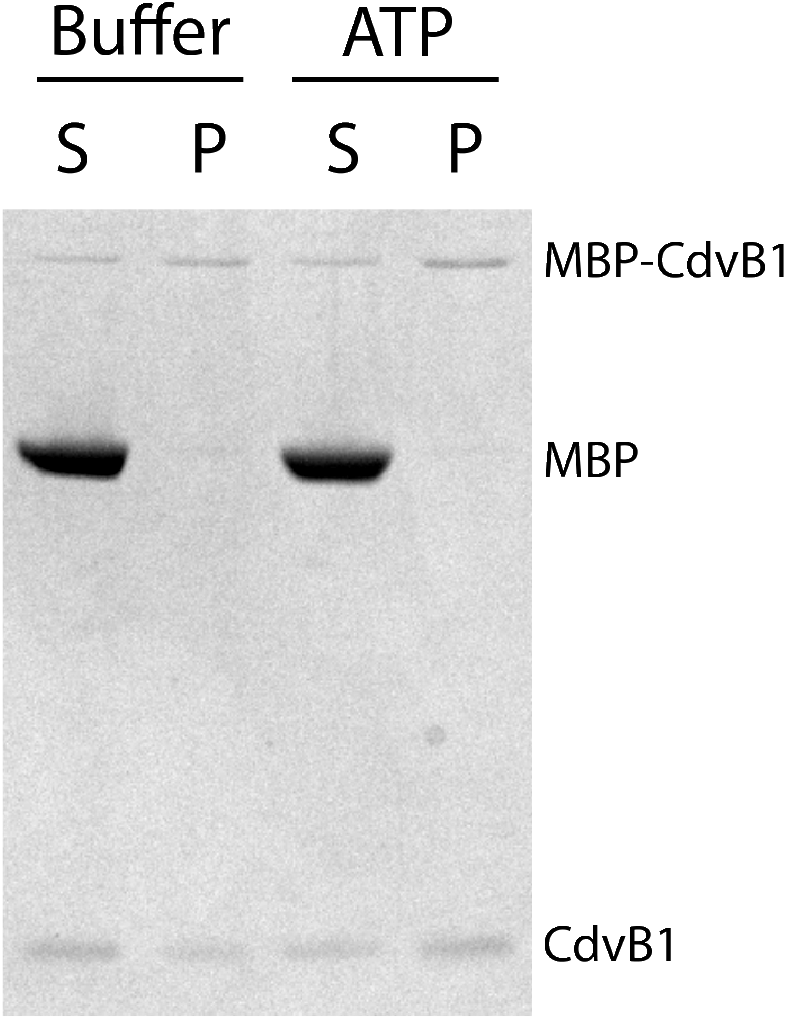
Pelleting assay performed to samples containing only CdvB1 incubated at 50 °C. No depolymerization of CdvB 1 filaments was visible

**Supplementary Figure 2.**
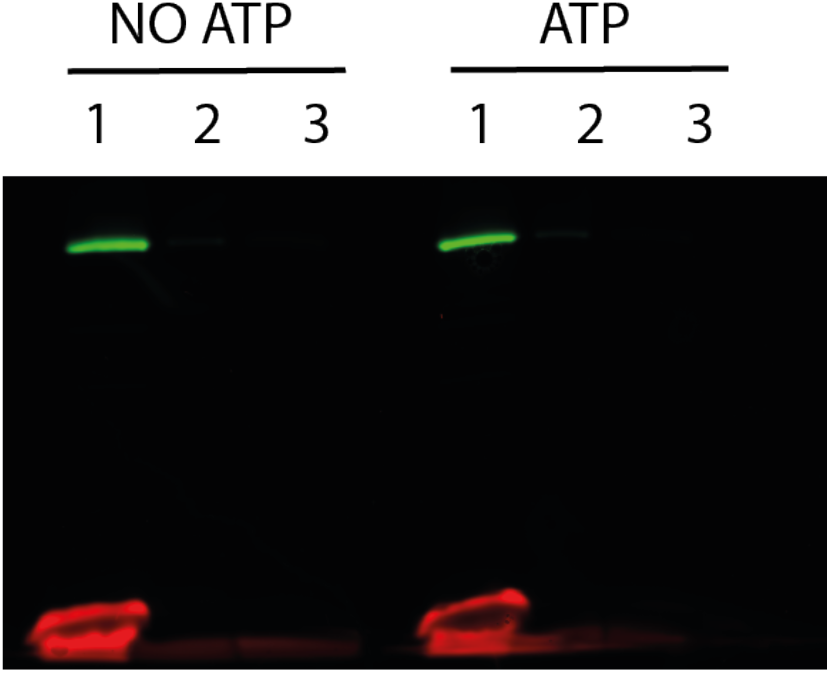
Membrane depolymerization control without any CdvC, where no depolymerization is visible in any case after the incubation at 50 °C with and without ATP

